# GeRelion: GPU-enhanced parallel implementation of single particle cryo-EM image processing

**DOI:** 10.1101/075887

**Authors:** Huayou Su, Wen Wen, Xiaoli Du, Xicheng Lu, Maofu Liao, Dongsheng Li

## Abstract

Single particle cryo-EM emerges as a powerful and versatile method to characterize the structure and function of macromolecules, revealing the structural details of critical molecular machinery inside the cells. RELION is a widely used EM image processing software, and most of the recently published single particle cryo-EM structures were generated by using RELION. Due to the massive computational loads and the growing demands for processing much larger cryo-EM data sets, there is a pressing need to speed up image processing. Here we present GeRelion (https://github.com/gpu-pdl¬nudt/GeRelion), an efficient parallel implementation of RELION on GPU system. In the performance tests using two cryo-EM data sets, GeRelion on 4 or 8 GPU cards outperformed RELION on 256 CPU cores, demonstrating dramatically improved speed and superb scalability. By greatly accelerating single particle cryo-EM structural analysis, GeRelion will facilitate both high resolution structure determination and dissection of mixed conformations of dynamic molecular machines.

---

Single particle cryo-EM has become a game-changing technique in structural biology, gaining unprecedented insights into many macromolecular machines in fundamental life processes^1,2^. By directly imaging the biological samples frozen in buffer solutions, cryo-EM circumvents the need to obtain well-ordered crystals, a major bottleneck in traditional crystallographic methods for high resolution structure determination. Single particle cryo-EM is also capable of computationally separating mixed conformations in a single sample, greatly facilitating the determination of high resolution structures and the analysis of dynamic molecular machines^3^.

In single particle cryo-EM, the molecules embedded in vitreous ice adopt different orientations, and generate various two dimensional (2D) projection images, so called particles. A three dimensional (3D) reconstruction is generated by averaging a large number of particles according to their rotation angles and in-plane shifts. The resolution of 3D reconstruction is gradually improved by iteratively refining the geometric parameters of each particle^4^. Furthermore, sophisticated computation methods enable the selection of the most homogeneous particles to achieve high-resolution 3D reconstructions, as well as the separation of different conformations within one sample. Particularly, RELION implements a Bayesian approach and the maximum a posteriori (MAP) algorithm, and demonstrates outstanding performance in 3D classification and 3D refinement^5,6^. In fact, most of the recently published single particle cryo-EM structures were generated using RELION. However, the computational cost of RELION is very high, and the enormous computational loads practically limit the number of cryo-EM particles that can be routinely analyzed. With the development of automatic EM data collection and the ever-growing demands for analyzing more particles to achieve higher resolution and to dissect mixed conformations, there is a pressing need to speed up the image processing^1,7^.

To accelerate the computation, many EM software packages, such as SPIDER^8^, FREALIGN^9^, EMAN^10^, and RELION^6^, implement parallel computation based on Central Processing Units (CPUs), and the speed increase depends on the number of available CPUs. In the last decade, due to the progress of hardware architecture and high-level programming model, the Graphic Processing Unit (GPU) has been widely used to accelerate time-consuming scientific applications such as medical image processing^11^, bioinformatics^12^ and machine learning^13^. Equipped with thousands of processing units in one processor, modern GPUs provide massive computation power, with significantly higher memory bandwidth than multi-core CPUs. In addition, the performance-price ratio of GPU is usually higher than that of CPU. Thus, GPU offers an attractive alternative for parallel implementation of the computation-intensive EM image processing applications. GPU implementation has been reported for EMAN, FREALIGN, and other programs^10,14,15^. Despite being extremely popular in the field for several years, the computationally expensive RELION has not been implemented in GPU system, probably due to the challenges to restructure the complicated work flow to map GPU computation. First, RELION uses coarse parallelization, which is not compatible with fine-grained parallelization of GPU. Second, the data structure in RELION is designed for individual images, resulting in discontinuous processing of multiple images. Third, RELION introduces two sampling methods, and the second sampling is sparse and sometimes with uncertainty, which makes it difficult for GPU implementation.

Here we present GeRelion, an efficient parallel implementation of RELION using GPU-enhanced system. Our tests on the two single particle cryo-EM data sets demonstrate that GeRelion with 4 or 8 NVIDIA GPUs outperforms RELION running on a modern CPU cluster with 256 CPU cores. GeRelion displays essentially linear scalability up to the 8 GPUs tested in our experiments, indicating great potential for further scaling on larger GPU clusters. In this paper we report the implementation of GeRelion, the results of performance tests, and a detailed procedure of utilizing the GeRelion program. This new parallel implementation on GPU system will significantly improve the efficiency of single particle EM image processing, and enable the routine analysis of much larger data sets, thus facilitating the determination of high resolution cryo-EM structures and the analysis of mixed conformations of dynamic molecular machines.

## RESULTS

### Profiling of RELION

RELION integrates various function modules for single particle cryo-EM image processing. Our implementation focuses on the two mostly used and computationally intensive functionalities: 3D classification and 3D refinement. In order to parallelize RELION on GPU, we first analyzed the RELION program to locate the computation hot spots and identified the computation patterns.

The mathematic basis of RELION is Bayesian statistics and the underlying algorithm is expectation-maximization method (details in **Online Methods**)^6^. We analyzed the execution time distribution of RELION, according to the program skeleton of RELION (**Supplementary Fig. 1**). The overall computation procedure is divided into three steps: expectation, maximization, and others. The expectation step consists of four major subroutines: getFourierTransformAndCtfs (getImg), getAllSquaredDifferences (getDiff), convertAllSquaredDifferencesToWeights (convert) and storeWeightedSums (store). The others step includes MPI communication, overhead of data read/write, and certain data processing tasks on the host side such as combining the partial 3D reconstruction files. We first used 8 CPU cores with MPI parallelism to evaluate the execution time distribution of RELION, by running 3D classification of the TRPV1 cryo-EM data set^16^. This data set contains 35,645 particles with a size of 256 x 256 pixels. The results show that the most time-consuming step is expectation, occupying over 97% of the total time, while the maximization and others steps take only 0.4% and 2.1%, respectively (**Fig. 1a**). Within the expectation step, the getDiff subroutine, which computes the l2-norm of expectation algorithm, is the slowest part. The same test was carried out for another single particle cryo-EM data set of the RAG complexes^17^, which contains 154,984 particles with a size of 192 x 192 pixels, resulting in a similar pattern of execution time distribution (**Fig. 1b**).

**Figure 1.**
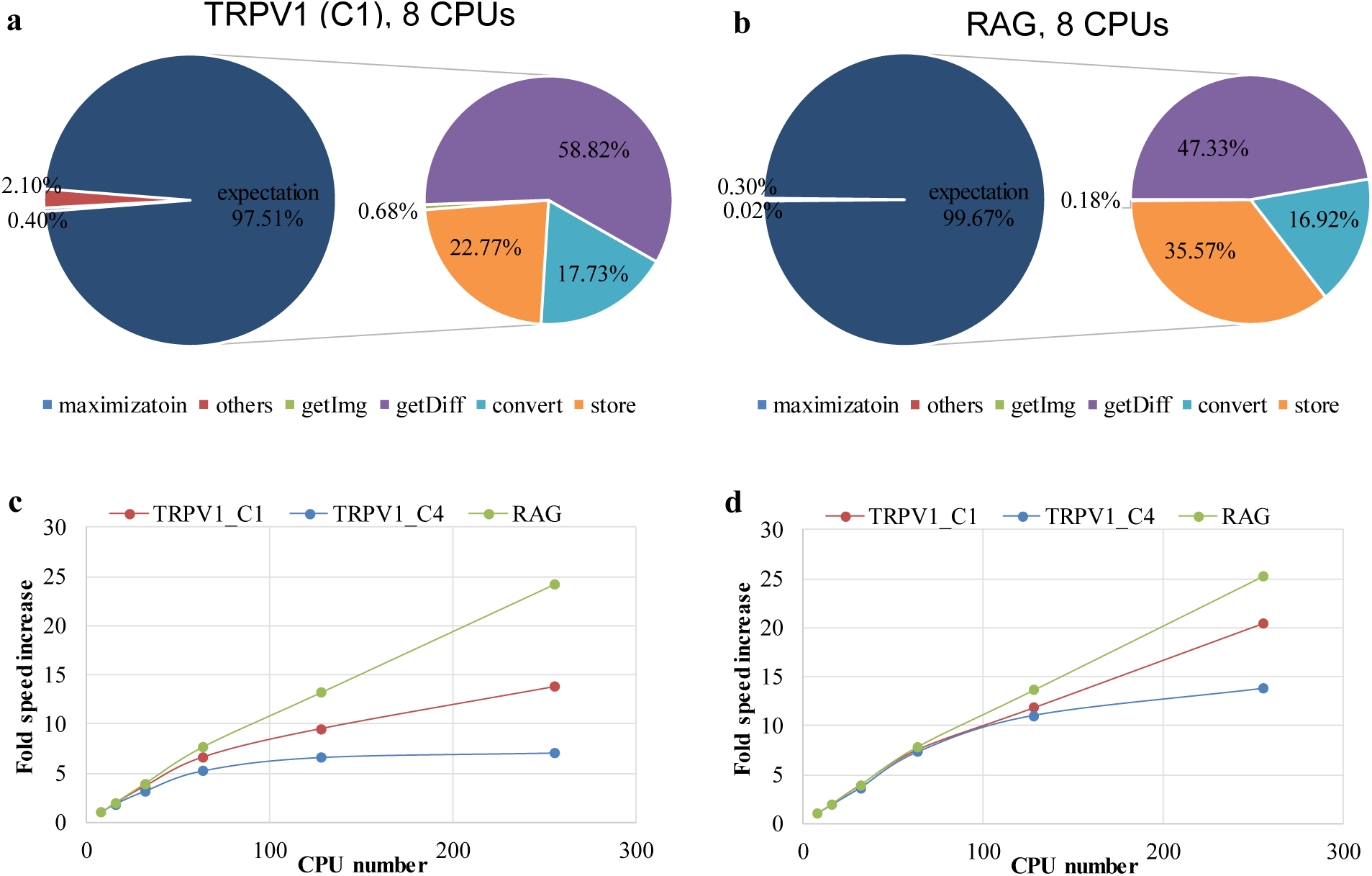
Execution time distribution and scalability of RELION. (**a**) Execution time distribution of 3D classification of the TRPV1 data set (without symmetry) using 8 CPU cores. The time distributions for the three steps (expectation, maximization and others) and for the four major subroutines (getImg, getDiff, convert and store) in the expectation step are shown. (**b**) Same as (**a**), except the 3D classification was carried out for the RAG data set. (**c**) Speed enhancement in 3D classifications of the TRPV1 data set, with and without C4 symmetry, and that of the RAG data set, using 8, 16, 32, 64, 128 and 256 CPU cores. The speeds of 8 CPU cores were used as the reference to calculate the speed enhancement. (**d**) Same as (**c**), except the speed enhancement is for the expectation step.

We further evaluated the scalability of RELION with increasing number of CPUs, by testing the total execution time and the time in the expectation step. We carried out 3D classifications using the TRPV1 data set with and without C4 symmetry, and also the RAG data set. The RELION computation speed on the TRPV1 data set increased linearly with up to 64 CPUs, but the speed improvement significantly slowed down with more than 64 CPUs (**Fig. 1c**). The limitation of CPU-based speedup was due to the substantially increased time proportions in the maximization and others steps (**Supplementary Fig. 2b**). The overhead of combining the partial 3D reconstruction files seems to be the most significant portion in the others step. The suppression of speedup with more CPUs appeared less severe in the tests on the TRPV1 data set without applying symmetry, and on the RAG data set that contains more than 4 times the particles of the TRPV1 data set (**Fig. 1c**). This is due to the increased computational loads in the slowest expectation step, which partially alleviated the increasing time occupancy in the maximization and others steps (**Supplementary Fig. 2e**). Indeed, for all three tests, the speedup in the expectation step was consistently more linear than that in the overall execution time (**Fig. 1d**). To remove the extra bottlenecks from the maximization step and certain overheads, which would significantly impact the scaling of RELION computation, we decided to parallelize all the steps on GPU system.

Based on the abovementioned tests and careful analysis of RELION program, we classified the RELION computation tasks into three categories: intensive computation, sparse index deduced computation, and global reduction operation. Specific strategies were developed for the parallel implementation of these different tasks on GPU, as detailed in **Online Methods**.

## Implementation of GeRelion

We have implemented GeRelion, a GPU-enhanced version of RELION, to accelerate the most widely used functionalities: 3D classification and 3D refinement (“auto-refine”). The original RELION codes for data read/write and MPI communication are unmodified, and the flow control of progressive processing in the original RELION is kept.

Rich data-level fine-grained parallelism and efficient memory access are two major factors that determine the performance of GPU program. In this work, we designed a four-level parallel model for the single particle cryo-EM image processing to exploit the powerful computational capability of GPU. In the first level, the particle images are divided into a set of pools that are parallelized onto individual GPUs, and the number of pools to be processed simultaneously equals to the number of GPUs (**Fig. 2a**). In the second level, the images within one pool are parallelized onto the stream multi processors of one GPU (**Fig. 2b**). In the third level, the workload for one image in one orientation will be assigned to one thread block of the GPU kernel, and the parallel degree of this level equals the number of orientations to be processed (**Fig. 2c**). In the fourth level, the processing of one or several pixels is assigned to one thread within each thread block, and all the pixels within one particle image are processed simultaneously (**Fig. 2d**). In GeRelion, the maximal parallel degree equals nr_gpus*M*N*K, where nr_gpus, M, N and K represent the number of GPUs, the number of images within one pool, the number of orientations to be processed, and the pixel number in one particle image, respectively.

**Figure 2.**
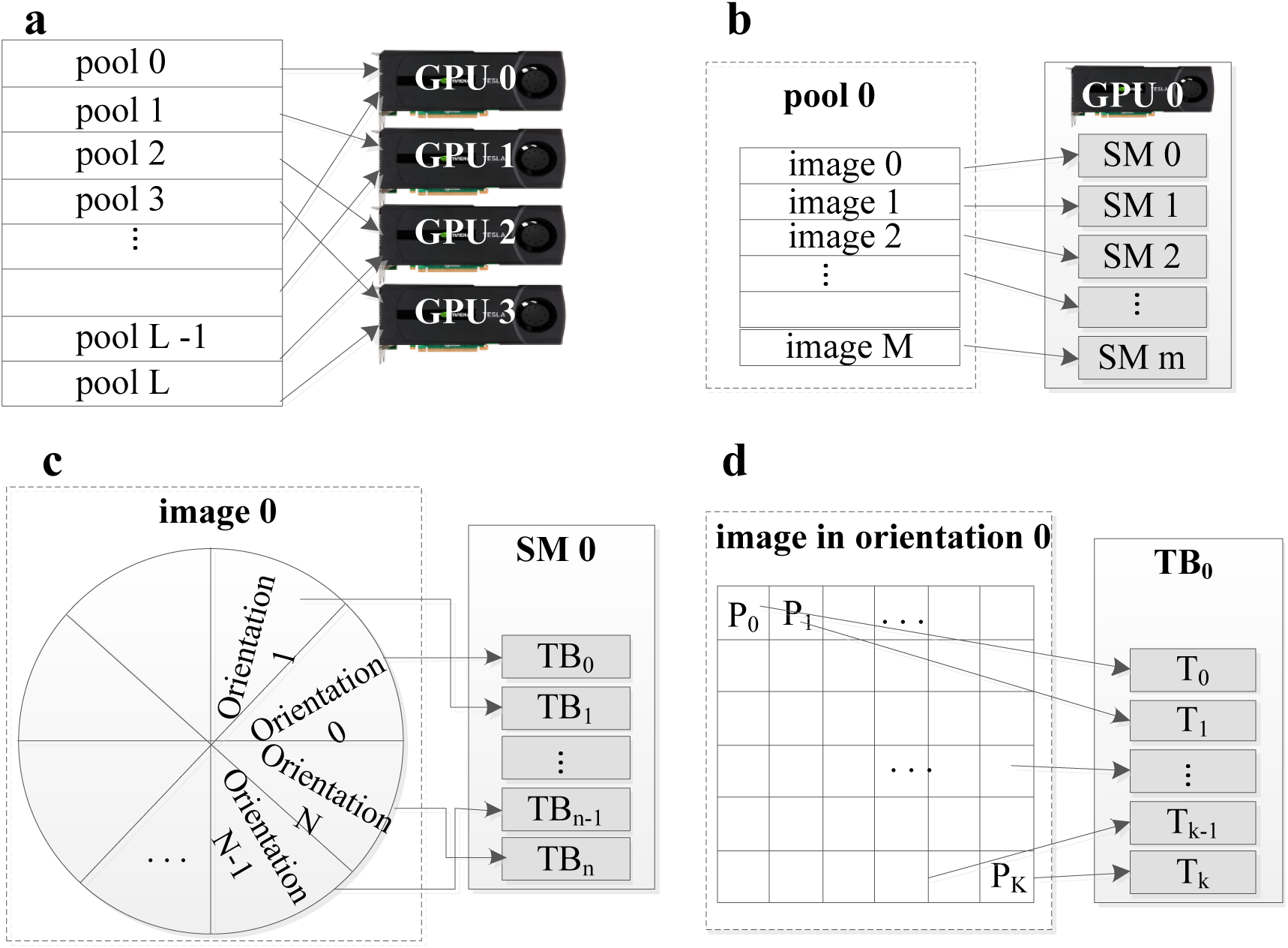
Multi-level parallel model of GeRelion. (**a**) Multi-pool parallelization to multiple GPUs. The particle images are divided into a set of pools. The maximal parallel degree is the number of GPU: nr_gpus. **(b)** Multi-image parallelization to stream multi processors (SM) of one GPU. The maximal parallel degree is the number of particle images within the pool: M. **(c)** Multi-orientation parallelization to thread blocks (TB). The maximal parallel degree is the number of orientations to be calculated for the particle image: N. **(d)** Multi-pixel parallelization to threads (T). The maximal parallel degree is the total pixel number within one particle image: K. Therefore, the maximal overall parallel degree of GeRelion equals nr_gpus*M*N*K.

In order to efficiently map the proposed multi-level parallel model to GPU-based system, we restructured the program in several aspects, as detailed in **Online Methods**. The original deep loops were flattened and partitioned, and data layout was reorganized to fit the architecture of GPU (**Supplementary Fig. 3**). To address the problem of memory limitation on GPU, GeRelion implements an adaptive parallel framework to determine the number of orientations to be processed simultaneously, based on the available memory space. To exploit the data reuse, we enlarged the parallel granularity in the first coarse sampling pass, and designed the lightweight kernels in the second fine sampling to achieve sufficient parallelism. In addition, we parallelized the sparse index computation by compacting the sparse data array to a continuous vector (**Supplementary Fig. 4**), and optimized the global reduction operation with atomic operation strategy.

## Performance of GeRelion

To test the performance of GeRelion and compare it with the original RELION, we used two computation systems (system 1 and system 2 in **Table 1**). The unmodified RELION ran on a CPU-only cluster with 256 CPU cores, while GeRelion ran on a GPU-based cluster with 2 nodes, each containing 4 GPUs and 12 CPU cores.

**Table 1.** Computation systems used for the tests.

| System 1: CPU cluster | System 2: GPU-based cluster |  |
| --- | --- | --- |
| CPU | CPU | GPU |
| Intel E5-2620 v2 | Intel E5-2620 v3 | NVIDIA K40m |
| 2.10GHz | 2.40GHz | 0.75GHz |
| 2x6 cores per node | 2x6 cores per node | 4 GPUs per node |
| DDR3 1866MHz | DDR4 2133MHz | GDDR 5 3004MHz |
| 128GB per nodes | 128 GB per node |  |
| 22 nodes | 2 nodes |  |

We first used the TRPV1 data set to run 3D refinement without or with C4 symmetry, and compared the computation times of RELION and GeRelion. The test results show that the overall computation speed of GeRelion on 4 GPUs is similar to that of RELION on 256 CPU cores, and GeRelion demonstrated a near-linear speedup with up to 8 GPUs, indicating excellent scalability of the implementation (**Fig. 3a, 3c**). The speed enhancement of GeRelion in the total execution time is consistent with that in the expectation step (**Fig. 3a, 3c**), which was in turn contributed by the similar speedup of all four major subroutines in the expectation step (**Fig. 3b, 3d**). We notice the extra speed increase in the overall execution compared to the expectation step (**Fig. 3a, 3c**), which is due to the significant speedup in the steps of maximization and others (**Table 2**, **Supplementary Fig. 5**). To prove the computation accuracy of our implementation, we also confirmed that the 3D reconstructions generated by RELION and GeRelion are essentially identical (**Fig. 3e, 3f**).

**Figure 3.**
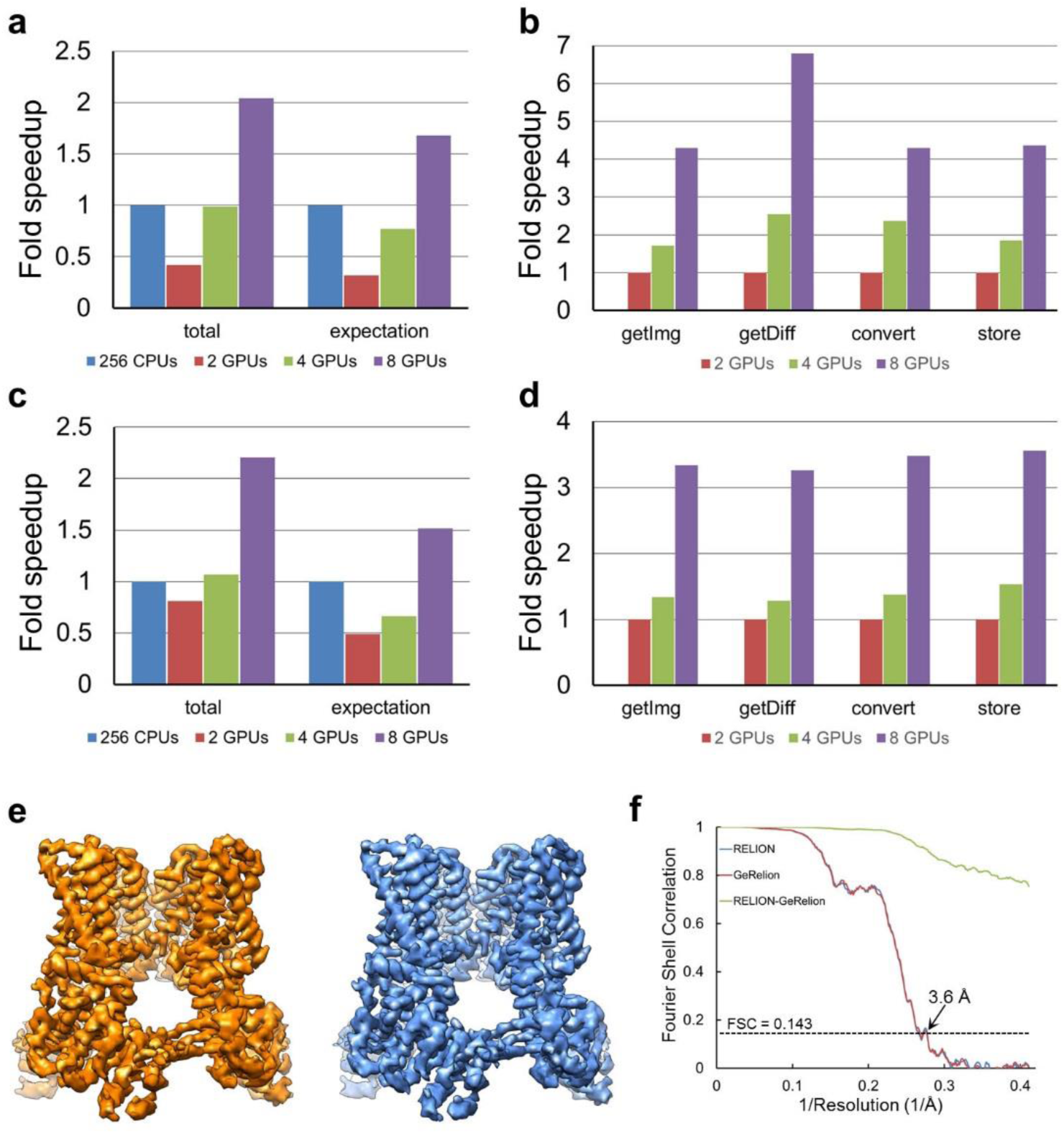
Performance of GeRelion in 3D refinement of the TRPV1 cryo-EM data set. (**a**) Computation speeds of RELION on 256 CPU cores and GeRelion on 2, 4 and 8 GPUs. The 3D refinement was carried out without applying symmetry. The speed of RELION was used as the reference to calculate the speed enhancement of GeRelion. (**b**) Computation speeds of the four major subroutines of the expectation step in GeRelion, from the tests in (**a**). The speeds of GeRelion on 2 GPUs were used as the reference to calculate the speed enhancement with more GPUs. (**c**) Same as (**a**), except the 3D refinement was carried out with C4 symmetry. (**d**) Same as (**b**), except the 3D refinement was carried out with C4 symmetry. All the corresponding execution times are listed in Table 2. (**e**) 3D reconstructions of TRPV1 generated by RELION (left, in orange) and GeRelion (right, in blue). Both maps were filtered at 3.6 Å resolution, and show essentially identical features. (**f**) Gold-standard Fourier shell correlation (FSC) curves (in orange and blue) of the two 3D reconstructions in (**e**) show excellent overlap, and the FSC curve between these two reconstructions (in green) shows the FSC values of 0.9 and 0.75 at the 3.6 Å resolution and Nyquist limit, respectively. These indicate that the two reconstructions generated by RELION and GeRelion are essentially identical.

**Table 2.** Execution time (in minutes) of RELION and GeRelion in auto-refine and 3D classification of the two single particle cryo-EM data sets.

|  |  | total | expecta-<br>tion | Maxi<br>mi-<br>zation | others | get-Img | getDiff | convert | store |
| --- | --- | --- | --- | --- | --- | --- | --- | --- | --- |
| TRPV1<br>auto-refine<br>C1 | 256 cpus | 298.0 | 221.2 | 50.0 | 26.8 | 1.0 | 164.2 | 10.0 | 26.3 |
|  | 2 gpus | 713.0 | 698.3 | 7.0 | 7.1 | 51.2 | 440.8 | 67.7 | 129.3 |
|  | 4 gpus | 301.0 | 287.4 | 7.7 | 5.9 | 29.9 | 173.3 | 28.6 | 70.1 |
|  | 8 gpus | 146.0 | 131.6 | 8.8 | 5.6 | 11.9 | 64.9 | 15.7 | 29.6 |
| TRPV1<br>auto-refine<br>C4 | 256 cpus | 203.0 | 115.9 | 60.4 | 26.7 | 0.9 | 82.8 | 3.4 | 16.4 |
|  | 2 gpus | 251.0 | 237.3 | 8.1 | 5.7 | 35.9 | 102.5 | 25.4 | 65.6 |
|  | 4 gpus | 190.0 | 174.6 | 8.5 | 7.3 | 26.9 | 80.0 | 18.5 | 43.0 |
|  | 8 gpus | 92.0 | 76.4 | 9.4 | 6.2 | 10.8 | 31.4 | 7.3 | 18.5 |
| RAG<br>auto-refine | 256 cpus | 863.1 | 773.0 | 69.0 | 21.0 | 34.2 | 517.7 | 82.5 | 203.2 |
|  | 2 gpus | 1697.0 | 1670.3 | 4.5 | 22.3 | 147.6 | 619.8 | 193.0 | 676.6 |
|  | 4 gpus | 1112.0 | 1071.9 | 4.5 | 22.7 | 110.6 | 470.0 | 140.4 | 304.7 |
|  | 8 gpus | 639.0 | 615.8 | 3.1 | 20.2 | 57.2 | 248.4 | 72.3 | 203.8 |
| TRPV1<br>3D<br>class<br>C1 | 256 cpus | 223.0 | 136.0 | 21.2 | 65.8 | 0.6 | 82.8 | 16.0 | 18.7 |
|  | 2 gpus | 314.0 | 302.3 | 1.3 | 10.5 | 9.6 | 160.4 | 63.3 | 64.7 |
|  | 4 gpus | 165.0 | 153.5 | 0.8 | 10.7 | 5.0 | 80.7 | 32.2 | 33.3 |
|  | 8 gpus | 107.0 | 84.0 | 6.5 | 16.5 | 2.6 | 40.7 | 16.2 | 17.1 |
| TRPV1<br>3D<br>class<br>C4 | 256 cpus | 157.0 | 66.7 | 20.0 | 70.3 | 0.7 | 25.9 | 4.1 | 23.3 |
|  | 2 gpus | 161.0 | 143.2 | 1.9 | 15.8 | 10.1 | 56.7 | 31.1 | 41.0 |
|  | 4 gpus | 93.0 | 78.8 | 1.0 | 13.2 | 5.7 | 31.0 | 16.6 | 23.0 |
|  | 8 gpus | 75.0 | 46.3 | 6.9 | 21.8 | 2.8 | 15.5 | 8.2 | 11.5 |
| RAG<br>3D<br>class | 256 cpus | 876.0 | 748.8 | 30.9 | 96.3 | 1.8 | 427.5 | 141.8 | 127.0 |
|  | 2 gpus | 2482.0 | 2456.4 | 1.8 | 23.9 | 32.3 | 1022.4 | 488.5 | 844.2 |
|  | 4 gpus | 1285.3 | 1248.0 | 1.3 | 36.1 | 50.0 | 521.9 | 250.2 | 425.1 |
|  | 8 gpus | 650.7 | 623.4 | 2.2 | 25.2 | 24.4 | 260.9 | 125.3 | 214.9 |

“Total” indicates the total execution time of the whole application, which includes three steps: “expectation”, “maximization” and “others”. The “expectation” step consists of four major subroutines: “getImg”, “getDiff”, “convert” and “store”. The computation tasks in the “others” step include MPI communication, I/O overhead, and certain data processing tasks on the host side such as combining the partial 3D reconstruction files.

We then compared the performance of GeRelion and RELION in 3D classification. The results are very similar to those from the tests of 3D refinement, showing excellent scalability of GeRelion with increasing number of GPUs (**Fig. 4**). For the TRPV1 data set, GeRelion on 4 GPUs outperformed RELION on 256 CPU cores (**Fig. 4a**). Since the scalability of CPU-based RELION is improved with higher computational loads in the expectation step (**Fig. 1c, 1d**), for the RAG data set with significantly more particles, 8 GPUs were needed for GeRelion to outrun the 256 CPU core-powered RELION (**Fig. 4c**). In addition, we also tested the 3D refinement using the RAG data set and the 3D classification using the TRPV1 data set with C4 symmetry. The execution times of all the tests in this work are summarized in **Table 2**. Collectively, our extensive performance tests demonstrate the superb acceleration and scalability of GeRelion.

**Figure 4.**
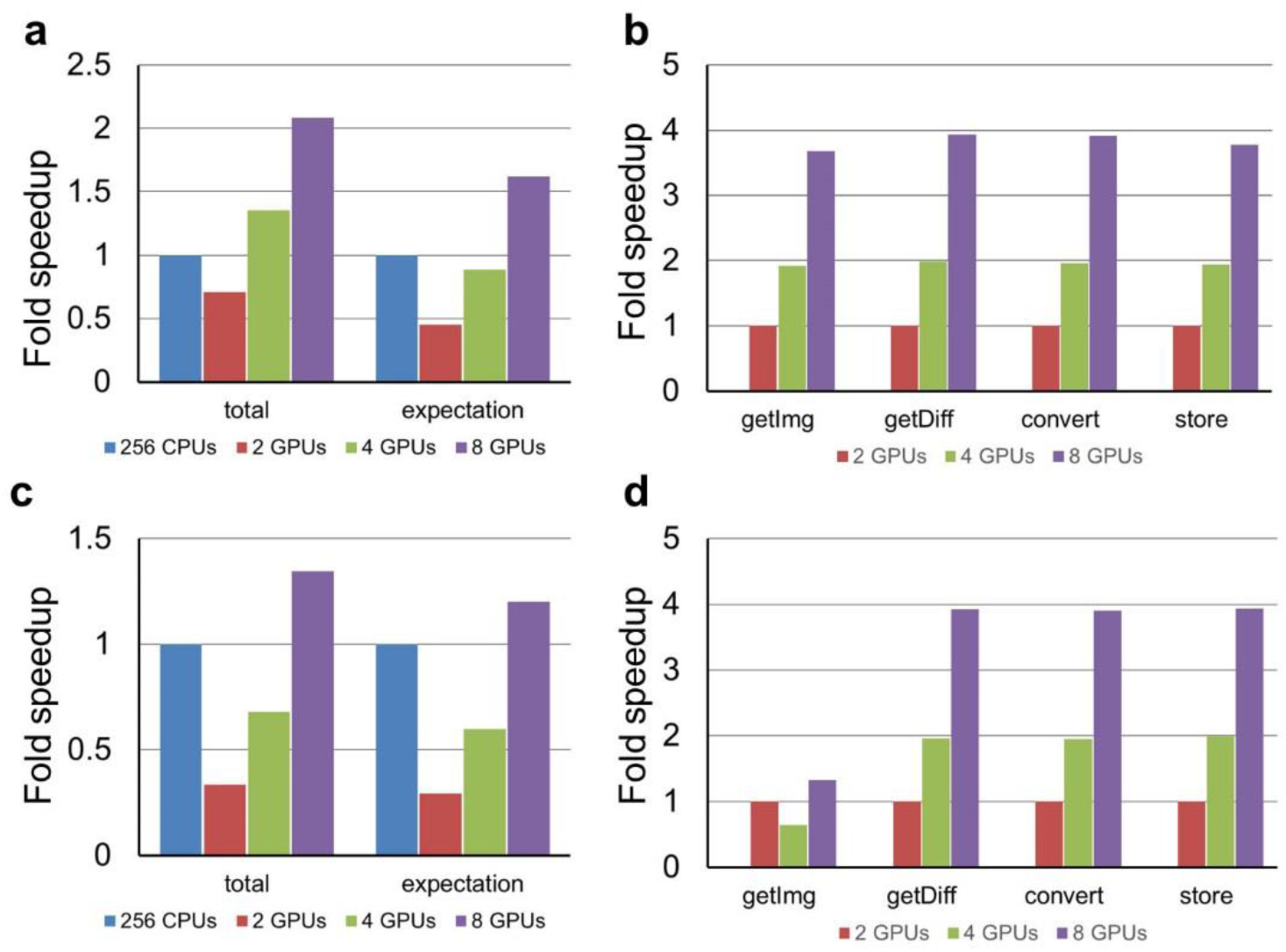
Performance of GeRelion in 3D classification. (**a**) Computation speeds of RELION on 256 CPU cores and GeRelion on 2, 4 and 8 GPUs, in 3D classification of the TRPV1 data set without symmetry. The speed of RELION was used as the reference to calculate the speed enhancement of GeRelion. (**b**) Computation speeds of the four major subroutines of the expectation step in GeRelion, from the tests in (**a**). The speeds of GeRelion on 2 GPUs were used as the reference to calculate the speed enhancement with more GPUs. (**c**) Computation speeds of RELION on 256 CPU cores and GeRelion on 2, 4 and 8 GPUs, in 3D classification of the RAG data set. The speed of RELION was used as the reference to calculate the speed enhancement of GeRelion. (**d**) Computation speeds of the four major subroutines of the expectation step in GeRelion, from the tests in (**c**). The speeds of GeRelion on 2 GPUs were used as the reference to calculate the speed enhancement with more GPUs. All the corresponding execution times are listed in **Table 2**.

## DISCUSSION

RELION is a popular image processing program in the cryo-EM field, and has generated most of the recently published high-resolution single particle cryo-EM structures. However, the Bayesian statistics-based computation in RELION is very costly, and the resulting massive computational loads limit the particle number of routine single particle data sets to well below one million. Due to the widely used automatic data collection and the need to analyze more particles to extract more structural information from a particular biological sample, there is a pressing need to significantly accelerate the image processing programs such as RELION^1^.

Here we presented GeRelion, an efficient GPU-based parallel implementation of the RELION program. To exploit the powerful computational capability of GPU system, we designed a four-level parallel model and restructure the RELION program to efficiently map this parallel model to GPU system. The resulting GeRelion implementation demonstrates significant speedup in the tests with two cryo-EM data sets, and shows excellent scalability with increasing number of GPUs. To our knowledge, GeRelion represents the first systematic GPU conversion of all the computation steps in the two most widely used RELION functionalities: 3D classification and 3D refinement.

Our tests show that GeRelion using 4 or 8 GPUs outperforms RELION using 256 modern CPU cores. Thus, to achieve similar computation speed, it is more cost-effective to purchase and maintain a couple GPU nodes than a large CPU cluster. Specifically, one NVIDIA K40m GPU provides 1.43T FLOPS (float-point operations per second) for double precision float, and its price is about $3,000. In contrast, an Intel Xeon E5-2620 v2 CPU supports up to 100.8G FLOPS, and its price is about $400 per CPU. When considering the cost of memory and other components of a computer node, the prices for 8 GPUs (2 nodes) and 256 CPU cores (22 nodes) are about $34,000 and $67,700, respectively. Practically it is also much easier to build and maintain a couple GPU nodes than a large CPU cluster. Furthermore, regular RELION computation uses double precision float point data, which makes it necessary to use high-end GPUs. The calculation with single precision float point has been implemented in the RELION version 1.4. If the computation with single precision float point can satisfy the requirement of image processing at least under certain circumstances, it will be an optimal choice to use the consumer level GPU at the price range of $200-600, which is similar to the price of a six-core Intel Xeon CPU.

Several aspects of our current GeRelion implementation may be further improved in the future work. First, the parallel degree of the getImg subroutine can be increased. In the current implementation, we only parallelized one pool of images for each kernel invoked, and the default number of 8 is insufficient for GPU. Therefore, we can parallel hundreds or thousands of images in this step and keep the results in GPU memory for the following getDiff subroutine. Second, the maximization step can be optimized for the large particles such as those with a size of 512 or more pixels. Due to the GPU memory limitation, currently it is difficult to execute the Fourier transform function for the very large particles. Third, a hybrid GPU-CPU implementation can be developed to utilize the CPU computation capability, which comes with the GPU system, for further improvement of the computation efficiency. Fourth, the parallel read/write can be improved, which will become particularly important for scaling in much larger GPU clusters.

In summary, GeRelion dramatically speeds up the computation-intensive 3D classification and 3D refinement in RELION, and demonstrates a great potential for scaling the speed enhancement on larger GPU clusters. GeRelion will greatly accelerate the processing of much larger single particle cryo-EM data sets, and thereby facilitate high resolution structure determination as well as analysis of mixed conformations in biological samples.

## METHODS

Methods and any associated references are available in the online version of the paper.

## ACKNOWLEDGEMENTS

The authors are grateful to S. Scheres (MRC) for sharing the source code of RELION, and would like to thank X. Zhu (Southern Medical University) for connecting the two research groups to initiate this collaborative work. This work is sponsored in part by the National Basic Research Program of China (973) under Grant No. 2014CB340303, the National Natural Science Foundation of China under Grant No. 61222205 and No. 61502509, the Program for New Century Excellent Talents in University, and the Fok Ying-Tong Education Foundation under Grant No. 141066, and so on.

## AUTHOR CONTRIBUTIONS

M.L. and D.L. conceived the project. D.L. and X. L. supervised the project. H.S. and D.L. designed the overall development of GeRelion. H.S., W.W., X.D and D.L implemented the GPU program. H.S, W.W and M.L designed the test and analyzed the results. M.L, D.L and H.S. wrote the manuscript. All authors read and approved the final manuscript.

## COMPETING FINANCIAL INTERESTS

The authors declare no competing financial interests.

TRPV1 (C1), 8 CPUs

RAG, 8 CPUs

## ONLINE METHODS

### RELION algorithm

RELION is an open-source software package for single particle cryo-EM structure determination. Like most of the existing implementations for cryo-EM image processing, it employs the so-called weak-phase object approximation, leading to the following linear image formation model in Fourier space.

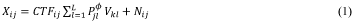

Where *X^ij^* is the *j*th component, with *j* = 1, …, J of the 2D Fourier transformation *X_i_* of the *i*th experimental image, with *i* = 1, …, N. *CTF^ij^* is the *j*th component of the contrast transfer function for the *i*th image. *V_kl_* is the *l*th component, with *l* = 1, …, L, of the 3D Fourier transformation *V_k_* of the *k*th of *K* underlying structures in the data set. All components *V_kl_* are assumed to be independent, zero-mean, and Gaussian distributed with variance 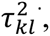 *P^Φ^* is a J × L projecting matrix of elements 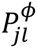, *N_ij_* is noise in the complex plane, which is assumed to be independent, zero-mean, and Gaussian distributed with variance 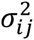.

The objection of cryo-EM image analysis software RELION is to find the model with parameter set Θ (including all 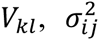, and 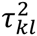) that has the highest probability of being the correct one in the light of both the observed data X and the prior information Y. From the Bayes’ law, this so-called posterior distribution factorizes into two components, shown as equation (2). Where the likelihood P(X|Θ, Y) quantifies the probability of observing the data given the model, and the prior P(Θ|Y) expresses how likely that model is given the prior information [16]. It should be noticed that most of the other methods in Fourier domain aimed to only optimize P(X| Θ, Y).

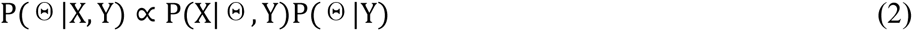

As for RELION, prior information is the smoothness hypothesis about cryo-EM reconstructions, and posterior information is provided by cryo-EM image data set. The deduced optimization is called maximum a posterior (MAP) estimation. MAP estimation is the most likely model which gets most out of the data at the Gaussian prerequisite of smoothness. RELION implements an iterative expectation-maximization algorithm to optimize the MAP model 2. The iterative algorithm is expressed by formulas 3∼7.

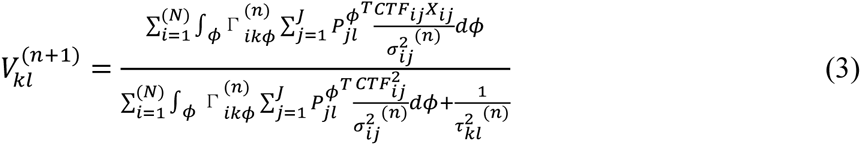

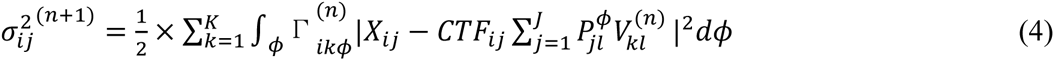

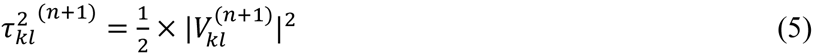

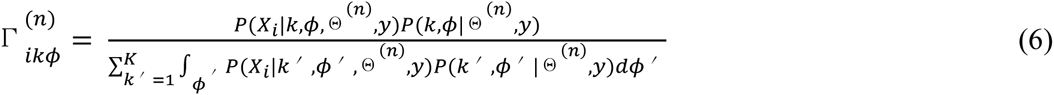

where 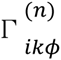 in equation (6) is the posterior probability of class assignment *k* and orientation assignment *φ* for the *i*th image, given the model at iteration n. The iterative algorithm starts from an initial estimate model of *V*_*k*_. User controls the number of models *K* that is to be refined simultaneously. Initial estimations for 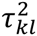 and 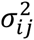 are calculated from the power spectra of the starting model and individual particles, respectively. Within each iteration, the posterior probability 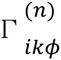 for all images has to be calculated in all possible orientation and class *k*. The major computation comes from the calculation of the *l*_2_-norm in equation (7).

### Three computation modes of RELION

According to the computation instructions, we classified the computation tasks of RELION into three categories: the intensive computation, the sparse index computation and the global reduction (**Supplementary Table 1**). In the first sampling of “getDiff”, the l2-norm will be calculated for all particle images and reference CTF images in all possible orientations. High resolution structure determination often requires hundreds of thousands, or even millions of particles, and more than a thousand of orientations for the angle search. On top of these, there is another search grid of XY shift. The total computation requirement for “getDiff” will be over PetaFLOPs, the procedure can be considered as computation intensive. In Relion, the process of the second fine sampling is only carried out in the significant orientations determined in the first sampling. An orientation can be consider as significant if the corresponding weight is larger than the threshold value. Compared with the total number of orientations, the number of significant orientations is relative small. The computation pattern is sparse index computation. To calculate weighted sum in the subroutine “store”, the operation of summing all images in formula (3) and the integration 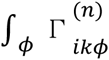 corresponds to solve the discrete sum in all significant orientations. Both of them refer to the global reduction for all particles and orientations. Similarly, the execution path of “store” also depends on whether the corresponding weight is significant. Due to the sparse feature of weight matrix, the process of “store” is both sparse index and global reduction.

### Restructuring of RELION for GPU implementation

In order to efficiently parallelize the RELION computation onto GPU system, we restructured the program in several aspects.

### Partition and unroll loops

The philosophy of GPU programing is unrolling loops to kernels. Generally, a simple kernel is designed based on one or several loops of code segment with the same parallel degree: the size of the loop index. In RELION, the subroutines consist of several layer of loops, the workload within different layer loops correspond to different parallel degrees. For example, the subroutine “getDiff” consists of three layer for loops, the orientation for loop, particle for loop and the translation for loop. The sub-functions of getFref, applyCTFtoFref and calDiff reside in these three layer loops respectively. That means the parallel degrees of them are not the same. These workloads should be parallelized with different GPU kernels. Therefore, we partition the long loops into several short loops, with the key function processed within each loop. Though loop unrolling, we can design kernels for each short loop by mapping workload of different layer loops to multilevel programming model of GPU (**Supplementary Fig. 3)**.

### Reorganize the data layout

In the original program, the image data of different particles are stored in separate memory space, which disobeys the continuous access principle of GPU programming. Additionally, in order to save memory space, the operation on-the-fly is adopted for projection and back-projection. In GeRelion, we reorganize the image data by gathering the raw image data of different particles into a large array at the beginning of each iteration. While in the following steps, all input data are generated from the previous kernels on GPU, which avoids the preparation of input for kernels and exploits the locality between producer and consumer.

### Build adaptive parallel framework

Generally, we hope to unroll the entire loop of a function to kernel. In RELION, in order to save memory, the program is designed to traverse loops to process all jobs of the entire orientation. Within each traversing, only one image will be processed. We consider to parallelize the workload of all orientations into one kernel, however, which may lead to out-of-memory error due to numerous orientations and images. In GeRelion, we proposed an adaptive parallel framework to address this problem. We firstly evaluate the maximal memory requirement according to the number of orientations, images and image size. Combining the memory requirement and the free GPU memory, a suitable parallel degree taken orientation as unit can be computed. Through this way, we can use the GPU resource as much as possible and avoid the problem of memory limitation.

### Enlarge parallel granularity and increase parallel degree

According to the profiling of RELION, there are three major computation modes. In GeRelion, we design kernels for different sampling passes. For the first pass, considering that most of the orientations is valid and there is good data locality between the whole XY shift grid, we design a coarse kernel to parallelize the first pass of getDiff by enlarge the parallel granularity. One thread is used to deal with the job of the whole XY shift grid, and the reference image data can be reused for multiple times. For the second sampling, where the significant orientations are very few, in order to exploit enough parallelism, we developed a lightweight kernel by giving priority to parallel degree and using one thread to process only one pixel.

### Transform sparse computation into continuous processes

Although the sparse index computation is difficult and inefficient for GPU, the GPU-based parallelization of the sparse computing is inevitable for achieving high-performance GPU enhanced RELION. On one hand, from the profiling results, the sparse computation occupies a certain proportion of the execution time, especially in 3D classification. On the other hand, if the sparse index computation functions remain on the host side, a lot of intermediate data must be copied back to CPU. However, the data transmission between CPU and GPU is costly. In GeRelion, we transform the sparse computation to continuous process to avoid divergence in GPU kernels. In our implementation, we pick up the significant weights of all particles from the sparse weight matrix into a small continuous vector. In order to keep the consistency of the weight with the corresponding image and orientation, an aux vector is introduced to store the indexes of the weights in the original sparse weight matrix. An example of the weight array and vector in CPU and GPU RELION are shown in **Supplementary Fig. 4**. Generally, the global reduction is done on the host side, even though the overhead of copy back the large weight matrix to CPU is costly and a lock primitive is also needed to ensure the correctness of the program. Due to the possible written conflict when back projecting 2D slices into the 3D Fourier space, we adopt atomic operation to implement global reduction.

### Performance tests of GeRelion

We used two single particle cryo-EM data sets to test the performance of GeRelion in 3D classification and 3D refinement (“auto-refine”). Both of the TRPV1 and RAG data sets can be downloaded from the Electron Microscopy Pilot Image Archive (https://www.ebi.ac.uk/pdbe/emdb/empiar/). The TRPV1 data set (EMPIAR-10005) contains 35,645 particles with a size of 256 x 256 pixels. For our tests, we only used the particles averaged from all 30 movie frames to generate a map at 3.6 Å resolution. A continued 3D refinement using the particles averaged from the #3-16 movie frames would have improved the resolution to 3.4 Å. The RAG data set (EMPIAR-10049) is a combination of SEC and PC particles, and contains 154,984 particles with a size of 192 x 192 pixels. In the tests of 3D classification, the TRPV1 data is classified into 3 classes, and the RAG data into 6 classes.

Currently GeRelion can only support command lines to submit jobs. The commands for running all the GeRelion tests in this work are listed below. “Node01” and “node02” are the hostnames of the two GPU nodes. “Relion_refine_mpi” is the executable file of GeRelion. GeRelion can run on GPU or CPU, by setting the “model” parameter to 1 or 0.

The command of auto-refine on TRPV1 without symmetry: mpirun --np 9 -N 5 --host node01,node02 relion_refine_mpi --o Refine_OPT_C1/run8 -¬auto_refine --split_random_halves --i new_DFMerge_20.star --particle_diameter 160 --angpix 1.2156 --ref EMD-5778.mrc --firstiter_cc --ini_high 60 --ctf --ctf_corrected_ref --flatten_solvent --zero_mask --oversampling 1 --healpix_order 2 --auto_local_healpix_order 4 --offset_range 5 -¬offset_step 2 --sym C1 --low_resol_join_halves 40 --norm --scale --j 1 --memory_per_thread 8 --dont_combine_weights_via_disc --mode 1

The command of auto-refine on TRPV1 with C4 symmetry: mpirun --np 9 -N 5 --host node01,node02 relion_refine_mpi --o Refine_OPT_C1/run8 -¬auto_refine --split_random_halves --i new_DFMerge_20.star --particle_diameter 160 --angpix 1.2156 --ref EMD-5778.mrc --firstiter_cc --ini_high 60 --ctf --ctf_corrected_ref --flatten_solvent --zero_mask --oversampling 1 --healpix_order 2 --auto_local_healpix_order 4 --offset_range 5 -¬offset_step 2 --sym C4 --low_resol_join_halves 40 --norm --scale --j 1 --memory_per_thread 8 --dont_combine_weights_via_disc --mode 1

The command of 3D classification on TRPV1 without symmetry: mpirun --np 9 -N 5 --host node01,node02 relion_refine_mpi --o Class3D_OPT/run8 --i allimg.star --particle_diameter 180 --angpix 1.23 --ref 3D_relion_class001_shz-5.dat -¬firstiter_cc --ini_high 40 --ctf --ctf_corrected_ref --iter 25 --tau2_fudge 4 --K 6 --flatten_solvent --zero_mask --oversampling 1 --healpix_order 2 --offset_range 5 --offset_step 2 --sym C1 -¬norm --scale --j 1 --memory_per_thread 8 --dont_combine_weights_via_disc --mode 1

The command of 3D classification on TRPV1 with C4 symmetry: mpirun --np 9 -N 5 --host node01,node02 relion_refine_mpi --o Class3D_OPT/run8 --i allimg.star --particle_diameter 180 --angpix 1.23 --ref 3D_relion_class001_shz-5.dat -¬firstiter_cc --ini_high 40 --ctf --ctf_corrected_ref --iter 25 --tau2_fudge 4 --K 6 --flatten_solvent --zero_mask --oversampling 1 --healpix_order 2 --offset_range 5 --offset_step 2 --sym C4 -¬norm --scale --j 1 --memory_per_thread 8 --dont_combine_weights_via_disc --mode 1

The command of auto-refine on RAG: mpirun --np 9 -N 5 --host node01,node02 relion_refine_mpi --o Refine3D_OPT/run8 -¬auto_refine --split_random_halves --i particles_autopick_sort_class2d.star --particle_diameter 200 --angpix 3.54 --ref 3i3e_lp50A.mrc --firstiter_cc --ini_high 60 --ctf --ctf_corrected_ref -¬flatten_solvent --zero_mask --oversampling 1 --healpix_order 2 --auto_local_healpix_order 4 -¬ offset_range 4 --offset_step 2 --sym D2 --low_resol_join_halves 40 --norm --scale --j 2 --memory_per_thread 4 --random_seed 1401784870 --dont_combine_weights_via_disc --mode 1

The command of 3D classification on RAG: mpirun --np 9 -N 5 --host node01,node02 relion_refine_mpi --o Class3D_OPT/run8 --i particles_autopick_sort_class2d.star --particle_diameter 200 --angpix 3.54 --ref 3i3e_lp50A.mrc --firstiter_cc --ini_high 50 --ctf --ctf_corrected_ref --iter 25 --tau2_fudge 2 --K 4 -¬flatten_solvent --zero_mask --oversampling 1 --healpix_order 2 --offset_range 3 --offset_step 2 ¬-sym C1 --norm --scale --j 1 --memory_per_thread 4 --dont_combine_weights_via_disc --mode 1

### Code availability

The GeRelion program is open source and available on github (https://github.com/gpu-pdl-nudt/GeRelion).

## References

1. Nogales, E. The development of cryo-EM into a mainstream structural biology technique. Nat Meth, 13, 24–27 (2016).

2. Bai, X., McMullan, G. & Scheres, S. H. . How cryo-EM is revolutionizing structural biology. Trends Biochem. Sci., 40, 49–57 (2014).

3. Liao, M., Cao, E., Julius, D. & Cheng, Y. Single particle electron cryo-microscopy of a mammalian ion channel. Curr. Opin. Struct. Biol. 27C, 1–7 (2014).

4. Cheng, Y., Grigorieff, N., Penczek, P. A. & Walz, T. A Primer to Single-Particle Cryo-Electron Microscopy. Cell, 161, 438–449 (2015).

5. Scheres, S. H. W. A Bayesian view on cryo-EM structure determination. J. Mol. Biol. 415, 406–18 (2012).

6. Scheres, S. H. W. RELION: Implementation of a Bayesian approach to cryo-EM structure determination. J. Struct. Biol. 180, 519–530 (2012).

7. Cheng, Y. Single-Particle Cryo-EM at Crystallographic Resolution. Cell, 161, 450–457 (2015).

8. Yang, C. et al. The parallelization of SPIDER on distributed-memory computers using MPI. J. Struct. Biol. 157, 240–249 (2007).

9. Grigorieff, N. FREALIGN: high-resolution refinement of single particle structures. J. Struct. Biol. 157, 117–25 (2007).

10. Tang, G. et al. EMAN2: an extensible image processing suite for electron microscopy. J. Struct. Biol. 157, 38–46 (2007).

11. Eklund, A., Dufort, P., Forsberg, D. & LaConte, S. M. Medical image processing on the GPU-Past, present and future. Med. Image Anal., 17, 1073–1094 (2013).

12. Luo, R. et al. SOAP3-dp: Fast, Accurate and Sensitive GPU-Based Short Read Aligner. PLoS One, 8, (2013).

13. Raina, R., Madhavan, A. & Ng, A. Y. Large-scale deep unsupervised learning using graphics processors. in Proc. 26th Annu. Int. Conf. Mach. Learn.-ICML’09 873–880 (ACM Press, 2009). doi:10.1145/1553374.1553486

14. Li, X., Grigorieff, N. & Cheng, Y. GPU-enabled FREALIGN: accelerating single particle 3D reconstruction and refinement in Fourier space on graphics processors. J. Struct. Biol. 172, 407–12 (2010).

15. Castaño-Díez, D. et al. Performance evaluation of image processing algorithms on the GPU. J. Struct. Biol. 164, 153–160 (2008).

16. Liao, M., Cao, E., Julius, D. & Cheng, Y. Structure of the TRPV1 ion channel determined by electron cryo-microscopy. Nature, 504, 107–112 (2013).

17. Ru, H. et al. Molecular Mechanism of V(D)J Recombination from Synaptic RAG1-RAG2 Complex Structures. Cell, 163, 1138–1152 (2015).

